# Trends and variation in Prescribing of Low-Priority Medicines Identified by NHS England: A Cross-Sectional Study and Interactive Data Tool in English Primary Care

**DOI:** 10.1101/211235

**Authors:** Alex J Walker, Helen J Curtis, Seb Bacon, Richard Croker, Ben Goldacre

## Abstract

**Background:** Routine accessible audit of prescribing data presents significant opportunities to identify cost-saving opportunities. NHS England recently announced a consultation seeking to discourage use of medicines it considers to be low-value. We set out to produce an interactive data resource to show savings in each NHS general practice, and to assess the current use of these medicines, their change in use over time, and the extent and reasons for variation in such prescribing.

**Results:** The total NHS spend on all low-value medicines identified by NHS England was £153.5m in the last year, across 5.8m prescriptions (mean £26 per prescription). Among individual medications, liothyronine had the highest prescribing cost at £29.6m, followed by trimipramine (£20.2m) and gluten-free foods (£18.7m). Over time, the overall total number of low-value prescriptions decreased, but the cost increased, although this varied greatly between medications. Annual practice level spending varied widely (median, £2,262 per thousand patients, IQR £1,439 to £3,298). The proportion of patients over 65 showed the strongest association with low-value prescribing; CCG was also strongly associated. Our interactive data tool was deployed to OpenPrescribing.net where monthly updated figures and graphs can be viewed.

**Conclusions:** Prescribing of low-value medications is extensive but varies widely by medication, geographic area and individual practice. Despite a fall in prescription numbers, the overall cost of prescribing for low-value items has risen. Prescribing behaviour is clustered by CCG, which may represent variation in medicines optimisation efficiency, or in some cases access inequality.

**Abbreviations:**

GP: General Practice
NHS: National Health Service
NICE: National Institute for Health and Care Excellence
PCA: Prescription cost Analysis
QOF: Quality Outcomes Framework

## Introduction

In July 2017 NHS England announced a list of 19 “low-value” treatments, which in their view should not routinely be prescribed in primary care, for wider consultation^1^. The stated goal of the consultation was to reduce unwarranted variation, provide clear guidance and reduce unnecessary spending^1^. The treatments listed were regarded as either ineffective, harmful, or very low-value. Examples include: glucosamine and chondroitin, which are prescribed for osteoarthritis associated pain, despite having only weak evidence of efficacy and being specifically recommended against by NICE^2^; and co-proxamol, a painkiller which was withdrawn in 2005 after safety concerns, but continues to be prescribed “off license” at very high cost.

In order to facilitate informed discussion of this topic, we aimed to summarise the current state of prescribing of these medicines in England in terms of overall spending, trends over time, regional variation, and individual practice-level variation. We also aimed to determine what practice level factors are associated with high prescribing of these low-value medicines. Lastly we set out to add accessible measures describing use of these low-value treatments to our widely used OpenPrescribing.net service, which provides open access to monthly data on treatments prescribed in each individual NHS general practice.

## Methods

### Study design

Our analysis is reported in compliance with the STROBE statement <u>(von Elm et al. 2014)</u>. It was a retrospective cohort study incorporating English GP practices, measuring variation in prescribing of low-value medications over time, geographically, and determining factors associated with the total cost of prescribing at the practice level. We use mixed effects linear regression to investigate correlation of total low-value prescribing cost with various practice characteristics. The low-priority medicines included are: Co-proxamol, dosulepin, doxazosin modified release, fentanyl immediate release, glucosamine and chondroitin, gluten free products, homeopathy, lidocaine plasters, liothyronine, lutein and antioxidants, omega-3 fatty acid compounds, oxycodone and naloxone combination product, perindopril arginine, rubefacients, tadalafil once daily, paracetamol and tramadol combination, travel vaccines, and trimipramine.

### Setting and data

We used data from our OpenPrescribing.net project, which imports prescribing data from the monthly data files published by NHS Digital^3^. These contain data on cost and volume prescribed for each drug, dose and preparation, for each month, for each English general practice. We identified all prescribing for each of the low-value medications in each month, and generated the composite prescribing measure described below. Each medication was identified using British National Formulary codes (see Appendix B for full code list). We also matched the prescribing data with publicly available data on practices from Public Health England^4^. This allowed us to stratify the analysis to look at reasons for variation in low-value prescribing at the practice level. Only standard English practices labelled within the data as a ‘GP practice’ were included within the analysis; this excluded prescribing in non-standard settings such as prisons. In addition, to exclude practices that are no longer active, those without a 2015/16 Quality Outcomes Framework (QOF) score and those with a list size under 1000 were excluded. Using largely inclusive criteria such as this reduced the likelihood of obtaining a biased sample.

### Total cost of low-priority prescribing

For each low-priority medicine, we calculated the total number of prescriptions, total cost and cost per prescription for the period July 2015 to June 2016, and July 2016 to June 2017. We used the ‘actual cost’ field rather than ‘net ingredient cost’; and calculated the change in prescription numbers, cost and cost per prescription between the two time periods.

### Low-priority measures and composite measure

We developed data queries to identify 18 of the treatments in the NHS Digital primary care prescribing data for England, as held in the OpenPrescribing database. We identified the drugs in the dataset using their BNF codes. Each measure was calculated as the total cost of low-priority items each month for each practice, divided by practice list size, to derive a prescribing rate of cost (GBP) per thousand patients. The NHS England advice on use of herbal remedies was not included in our analysis as it could not practically be turned into a series of BNF codes. We then generated a composite measure of all low-priority prescribing, which was defined as the total cost of all low-priority medicines for each practice, divided by each practice’s list size to produce a rate of cost per thousand patients. Where monthly data were not required, and in order to smooth prescribing rates over a year, we aggregated 12 months together to generate a rate of cost per thousand patient years.

### Geographical variation at CCG level

For each low-priority prescribing measure, rates were aggregated by grouping each practice to its parent Clinical Commissioning Group (CCG) and then described using a map in which each CCG’s prescribing was represented using a colour spectrum.

### Monthly trends and variation across practices

For each low-priority prescribing measure, we described the monthly trends between July 2012 and June 2017 by calculating deciles at practice level for each month and plotting these deciles. We also used a histogram to describe the distribution of low-priority prescribing volume amongst practices.

### Factors associated with low-value prescribing

We built a linear regression model to assess the extent to which variation in such prescribing was correlated with: composite QOF score, which is a performance management metric used for GPs within the NHS, produced by NHS Digital; practice list size (calculated as mean over the most recent year); Index of Multiple Deprivation (IMD) score; patients with a long term health condition (%); patients over 65 (%), and whether each practice is a ‘dispensing practice’ with an in-house pharmacy service (yes or no). We used the total cost of all low-priority prescribing per 1000 patients, aggregated over the previous year as the dependent variable.

We stratified the rate of low-priority prescribing according to the factors defined above. These factors were also entered into a linear regression model, then a mixed effects linear regression model with low-priority prescribing rate as the dependent variable, the above variables as fixed effect independent variables and the parent CCG of each practice as a random effect variable. Prescribing and other practice quality measures were divided *a priori* into quintiles for analysis, except for existing binary variables (i.e. dispensing practice status). This was done for ease of interpretation, and to allow for variables having a non-linear effect on prescribing cost. Practices with missing data for a particular variable were not included in models containing that variable. From the resulting model, incidence rate ratios were calculated, with corresponding 95% confidence intervals. The level of missing data was determined and reported for each variable.

### Interactive Data Tool

We imported all data onto OpenPrescribing.net and made a series of 19 standard measures, one for each of the medicines plus an ‘omnibus’ measure, which aggregates all the low-priority medicines together. Measures were calculated for each month as the total cost of the medicine divided by the population list size. Here we report on the subsequent web site access summary statistics and media coverage.

### Software and Reproducibility

Data management was performed using Google BigQuery and Python, with analysis carried out using Stata 13.1. Data and analytic code can be found here: https://figshare.com/s/1acc3609a854825cd930

## Results

In this paper we report on the aggregate of all low-priority prescribing combined, and illustrative examples of trends and variation from individual medications; our detailed report on trends and variation for prescribing of all individual medications in the NHS England consultation can be found in Appendix A. Individual CCG or practice level data can be explored further on the OpenPrescribing.net website, where all data is current and updated on a monthly cycle.

### Total expenditure for low-priority medicines

There was £153.5m of total expenditure across all low-priority medications between July 2016 and June 2017, from a total of 5.8m items (Table 1). This represents £2.63 per person per year (total list size population 58.3m), and around 1.7% of the overall NHS spend on primary care prescribing (£9.2bn in 2016)^5^. For individual medications, there was a large range of total spend for each medication, from £78k (homeopathy) to £29.6m (liothyronine). The overall cost per item for all medications was £26.28, which ranged from £2.55 (dosulepin) to £411.97 (liothyronine).

**Table 1.**
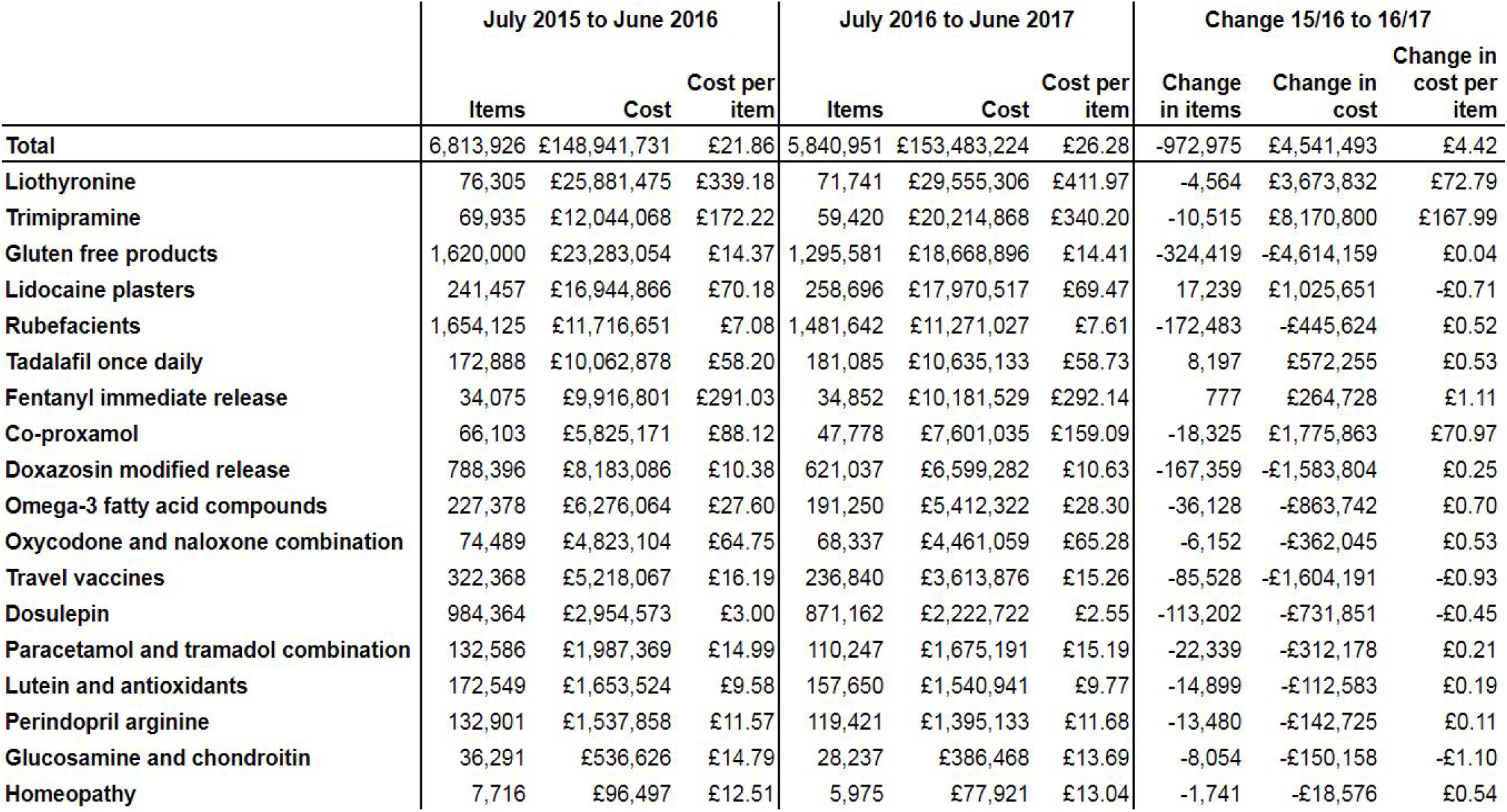
Total prescriptions, cost and cost per prescription in preceding two years for “low-priority” medications, along with change between 15/16 and 16/17 in English primary care.

### Trends in low-value prescribing

There has been an overall trend towards fewer low-value items being prescribed over time (Figure 1(a)), with almost one million fewer prescriptions (14% lower) between July 2016 and June 2017, compared to the previous year (Table 1). Despite the consistent downward trend in items, costs have risen, increasing by £4.5m (3% higher) when comparing 2016/17 with 2015/16 (Table 1). While the cost per item has remained stable for most medications, for liothyronine, trimipramine, and co-proxamol, the cost per item has risen dramatically (by £73, £168, and £71 per prescription respectively). In most individual practices, costs have risen (Figure 1(b)), although practices in higher deciles have risen more than those in lower deciles, where costs are relatively steady. For the individual measures, most are falling in prescribing volume, except for lidocaine plasters, tadalafil once daily and fentanyl immediate release, where small increases were observed between 2015/16 and 2016/17 (Table 1 and Appendix A). Some costs have fallen a great deal in this time period for (e.g. travel vaccines have fallen by 44%), but others have risen dramatically (e.g. trimipramine national costs have increased by 40%).

**Figure 1:**
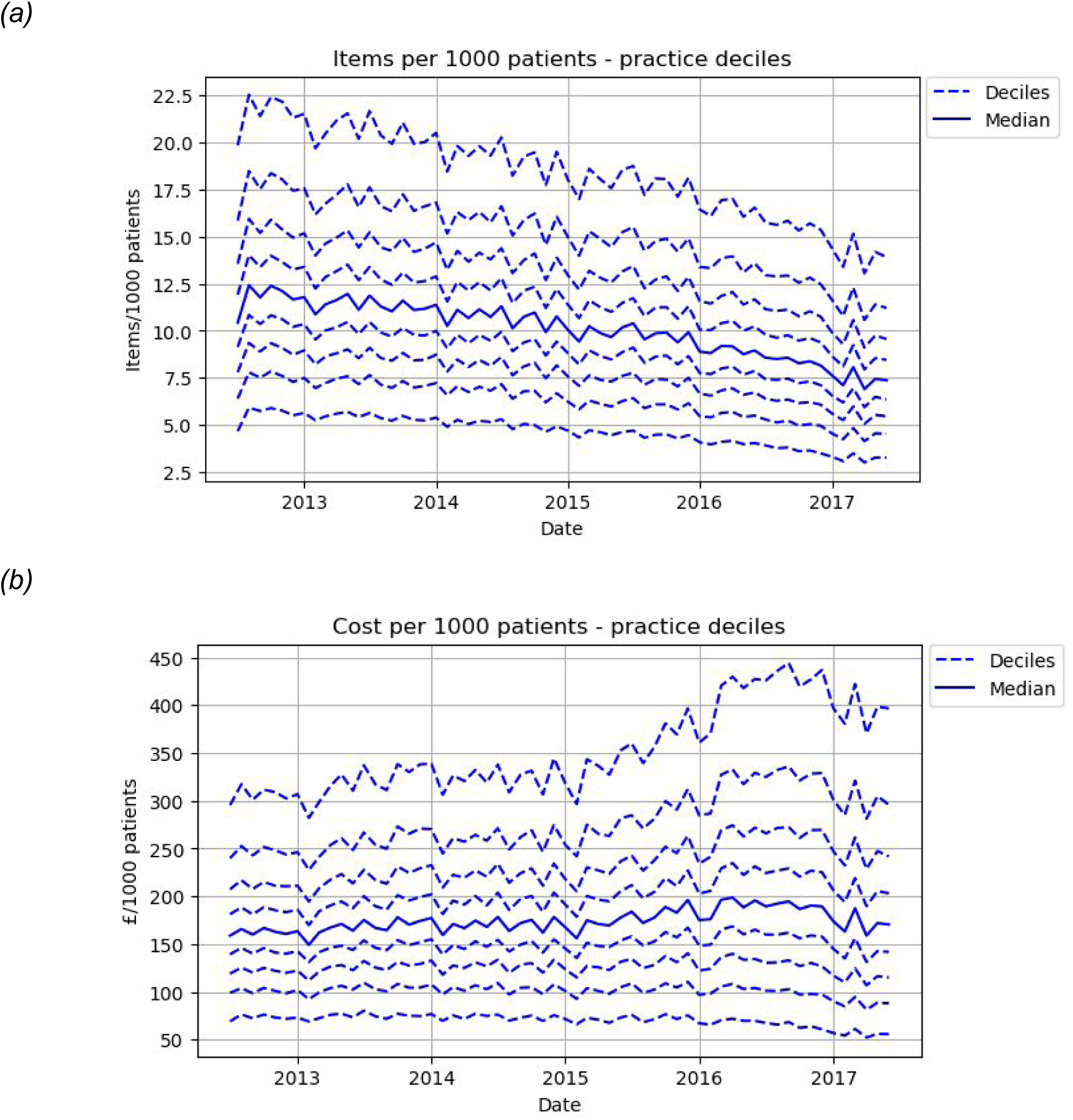
Total items (a) and cost (b) of prescribing for all low-value medicines combined over time in English primary care. Solid line is the median and surrounding dashed lines are deciles.

### Variation amongst practices and CCGs

The distribution of overall spending on low-priority medications is shown in Figure 2. Around half of practices are clustered within £1,000 of the median spend per thousand patients (median = £2,262, IQR £1,439 to £3,298). However, some practices spend very little per thousand patients, including 261 practices that spend less than £500 per thousand patients per year; while 588 practices spend more than £5,000 per thousand patients per year.

**Figure 2:**
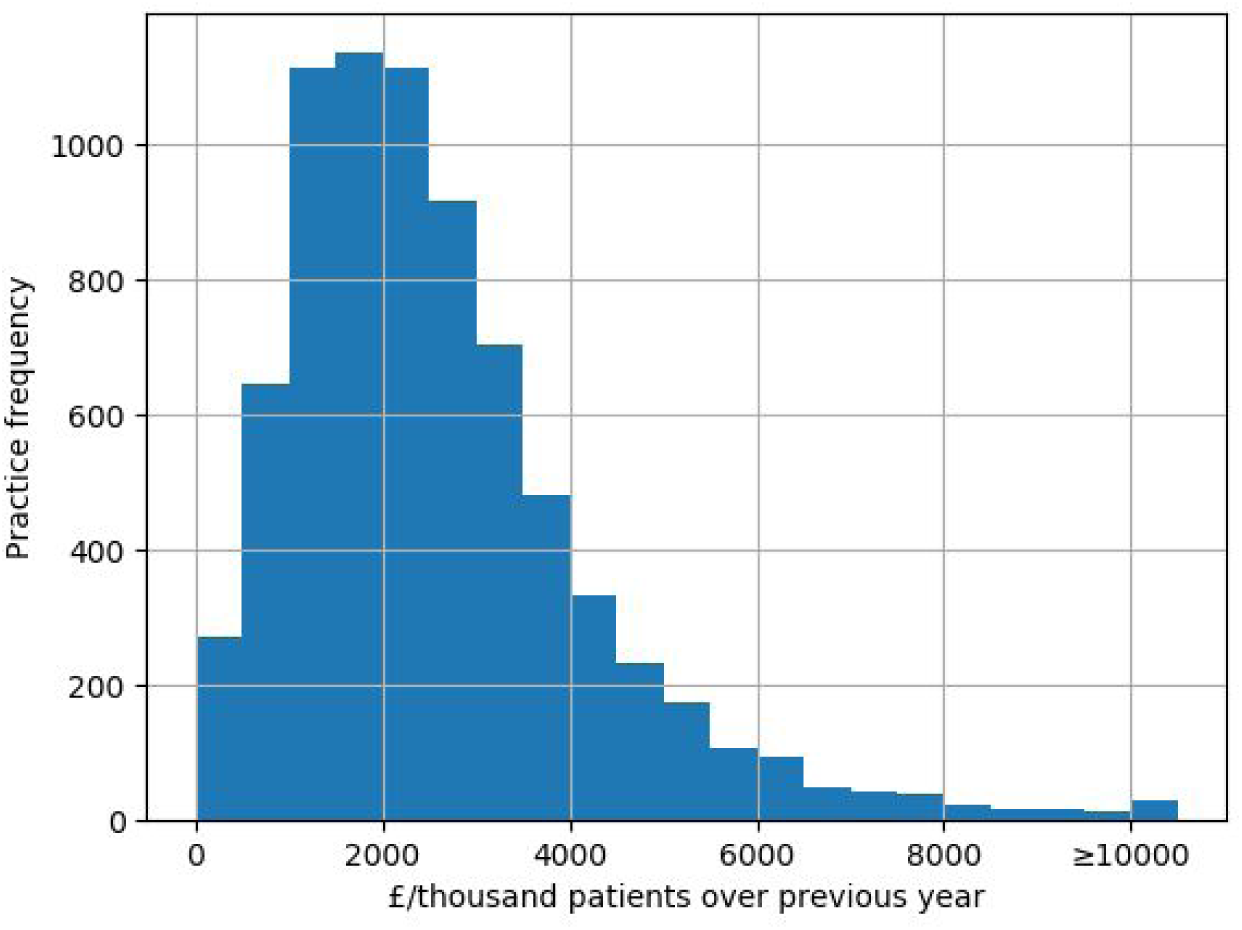
Distribution of total low-priority spending per practice, GBP per thousand patients per practice, between July 2016 and June 2017. Values over £10,000 are aggregated into the final column.

There are three medicines (liothyronine, trimipramine, and co-proxamol) where the volume of prescribing has decreased, but the total cost has dramatically increased (Table 1). This is shown in more detail in Figure 3 for co-proxamol, and in Appendix A for all individual medicines. Co-proxamol was withdrawn in 2005 and subsequently removed from the drug tariff. Prescribing has continued “off label”, in diminishing volumes, but the cost of the drug has risen dramatically over time: of note, was a shortage at the end of 2015 ^6^, hence the sudden drop in prescribing volume at this time, but with no reduction in linear downward trend in volume prescribed.

**Figure 3:**
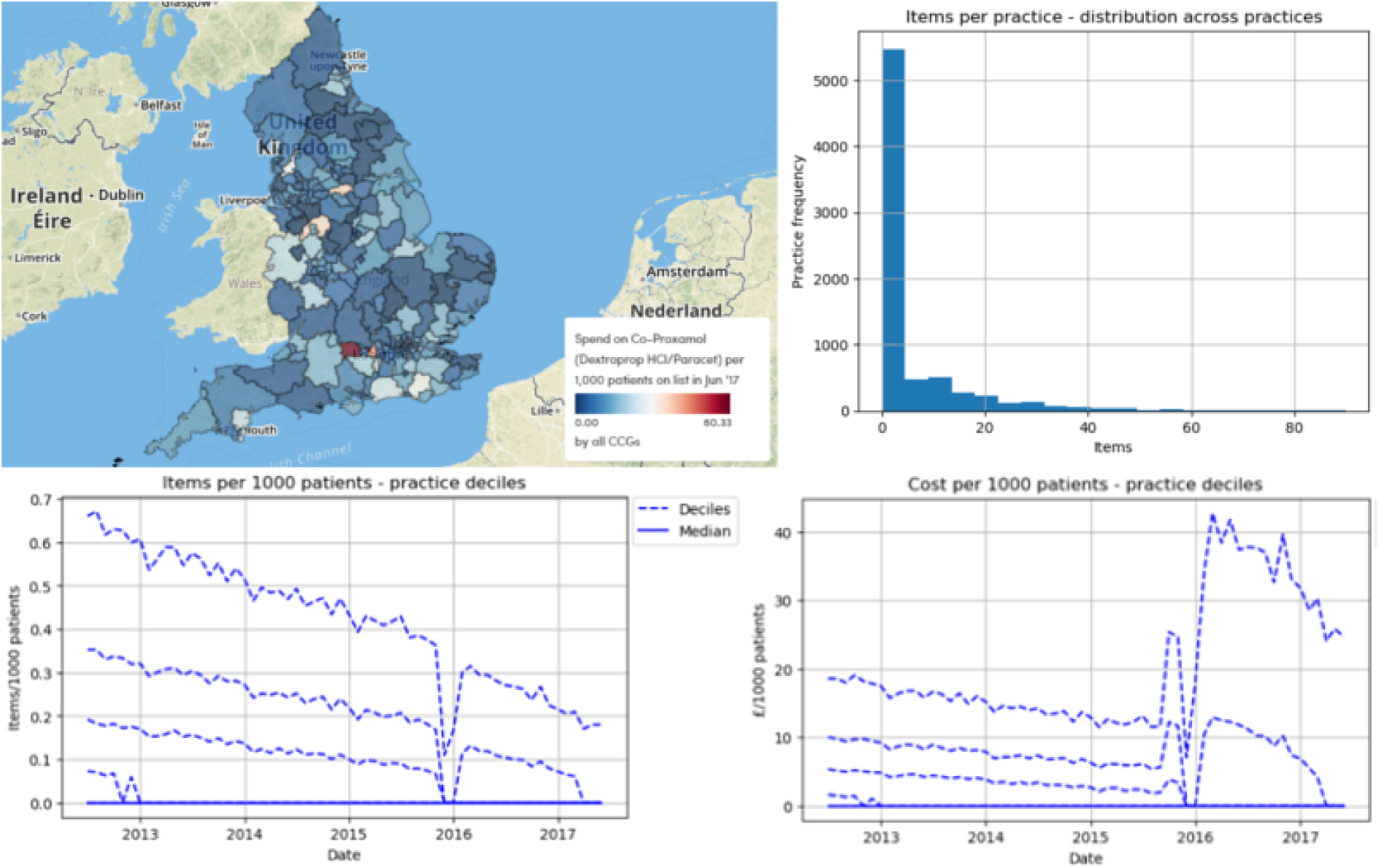
Variation in prescribing of co-proxamol, from top left to bottom right: variation in cost geographically at CCG level (£ per 1000 patients, June 2017); variation in items across practices within the last year; practice level variation in items over time; and practice level variation in cost over time.

Co-proxamol is also an example of a medication with significant costs to the NHS that is only prescribed by a small proportion of NHS practices. There are many other individual medications where over half of practices prescribe zero items per month, including: fentanyl immediate release, glucosamine and chondroitin, homeopathy, liothyronine, lutein and antioxidants, oxycodone and naloxone combination, paracetamol and tramadol combination, perindopril arginine, and trimipramine. Full details on prescribing of all these medications can be found in Appendix A.

### Factors associated with low-value prescribing spend

Overall, practices with a good QOF score spent slightly more per thousand patients than those with a poor score in univariate analysis (£165 more for the best score group vs the worst score 95% confidence interval £46 to £285). However, this effect is reversed in multivariable modeling, with only the ‘best’ vs ‘worst’ comparison being significant. Being in an area with a more deprived IMD score was associated with decreased spending in the univariable analysis, but not after multivariable adjustment. Practices with larger list sizes were associated with a slight increase in low-priority spending per thousand patients (£352 more per thousand patients in the largest practice group vs the smallest, multivariable CI: £238 to £467). Dispensing practices also spent slightly more (£255) per thousand patients than non-dispensing practices (multivariable CI: £133 to £377).

The percentage of patients over 65 had by far the strongest association with low-priority spending, of the factors assessed (Table 2). Practices with the highest percentage of patients over 65 (>22.5%) spent £1,302 more than those with the lowest percentage (<10.8%) after adjustment (multivariable CI: £1,139 to £1,466). While the percentage of patients with a long term health condition had a moderately strong association on low-priority spending in the univariable analysis, this effect was much reduced after multivariable modelling (Table 2). Within the mixed effects linear regression model, CCG as a random effect was found to be significantly associated with prescribing cost (p<0.0001), indicating clustering of spend on low-value medicines by CCG.

**Table 2.** Stratification and modelling of annual low-priority cost. Each independent variable is split into quintiles, linear regression modelling was performed to determine the change in cost associated with being in different quintiles of each variable.

|  | Quintile boundaries | Mean cost (£) per 1000 people | Change in cost (£)* | 95% confidence interval (£) |  |  | Multivariable change in cost (£)** | 95% confidence interval (£) |  |
| --- | --- | --- | --- | --- | --- | --- | --- | --- | --- |
| Quality outcomes framework (QOF) score (max = 559) | <522 | 2,494 | Reference |  |  |  | Reference |  |  |
|  | 522 to 541 | 2,553 | 58 | -62 | 178 |  | -33 | -143 | 76 |
|  | 541 to 550 | 2,578 | 84 | -36 | 203 |  | -87 | -197 | 24 |
|  | 550 to 556 | 2,655 | 160 | 41 | 280 |  | -53 | -164 | 59 |
|  | >556 | 2,660 | 165 | 46 | 285 |  | -156 | -271 | -41 |
| Index of Multiple Deprivation (IMD) | Least deprived | 2,859 | Reference |  |  |  | Reference |  |  |
|  | I | 2,754 | -105 | -225 | 14 |  | -108 | -224 | 9 |
|  | I | 2,579 | -280 | -400 | -161 |  | -125 | -253 | 2 |
|  | I | 2,413 | -446 | -565 | -327 |  | -150 | -290 | -9 |
|  | Most deprived | 2,337 | -522 | -641 | -403 |  | -114 | -274 | 46 |
| Practice list size | <3,784 | 2,340 | Reference |  |  |  | Reference |  |  |
|  | 3,785 to 5,796 | 2,531 | 191 | 72 | 311 |  | 220 | 112 | 329 |
|  | 5,798 to 8,018 | 2,666 | 326 | 207 | 446 |  | 279 | 169 | 389 |
|  | 8,020 to 11,165 | 2,747 | 408 | 288 | 527 |  | 364 | 252 | 476 |
|  | >11,165 | 2,656 | 316 | 197 | 436 |  | 352 | 238 | 467 |
| Dispensing practice? | No | 2,512 | Reference |  |  |  | Reference |  |  |
|  | Yes | 3,073 | 560 | 450 | 671 |  | 255 | 133 | 377 |
| % of patients over 65 | <10.8% | 1,788 | Reference |  |  |  | Reference |  |  |
|  | 10.8% to 15.4% | 2,411 | 623 | 508 | 738 |  | 556 | 438 | 674 |
|  | 15.4% to 18.8% | 2,642 | 854 | 739 | 969 |  | 801 | 669 | 933 |
|  | 18.8% to 22.4% | 2,910 | 1,121 | 1,006 | 1,236 |  | 1,040 | 897 | 1,184 |
|  | >22.4% | 3,191 | 1,402 | 1,287 | 1,517 |  | 1,302 | 1,139 | 1,466 |
| % of patients with a long term health condition | <47.0% | 2,113 | Reference |  |  |  | Reference |  |  |
|  | 47.0% to 51.5% | 2,557 | 444 | 326 | 563 |  | 194 | 82 | 307 |
|  | 51.5% to 55.3% | 2,680 | 567 | 448 | 686 |  | 221 | 106 | 337 |
|  | 55.4% to 59.7% | 2,757 | 645 | 526 | 763 |  | 165 | 44 | 285 |
|  | >59.7% | 2,839 | 726 | 607 | 844 |  | 216 | 89 | 342 |

\*linear regression
\*\*mixed effects linear regression, adjusted for: all other variables in table as fixed effects, plus CCG as a random effect

### Interactive Data Tool

The online data tool is now online on the OpenPrescribing web site ^7^ and can be explored at individual medication level, as well as practice and CCG level. The launch of the web site led to various news stories being published ^8–12^ and a good level of engagement with the site, with 1,286 pageviews (983 unique pageviews) between the launch date (2nd October 2017) and the most recent extraction of page view data (25th October 2017).

## Discussion

### Summary of findings

We found that prescribing of medications identified as low-priority by NHS England cost a total of £153.5m within the last year, or 1.7% of total NHS spending on primary care prescribing^5^. Despite a decrease in the total number of items prescribed over recent years, the total cost of these prescriptions has increased due to a large increase in the cost per prescription for some medications. There was wide variation in the total cost of prescribing amongst the identified medicines over the most recent year, from £29.6m for liothyronine to just £77,921 (and declining) for homeopathic products. The strongest association with the level of prescribing cost at practice level was with the proportion of patients over 65. This is perhaps not surprising given that older patients are generally more likely to receive prescriptions. Given the recent trends in medicines such as co-proxamol, glucosamine, homeopathy and trimipramine, it seems conceivable that prescribing of these medicines could approach zero within a few years. Our online interactive tool ^7^ will continue to provide useful and up to date information (updated monthly) on the continuing prescribing of these medications.

### Strengths and weaknesses

We included all typical practices in England, thus minimising the potential for obtaining a biased sample. We used real prescribing and spending data which are sourced from pharmacy claims and therefore did not need to rely on the use of surrogate measures. Using this data rather than survey data, also eliminates the possibility of recall bias. We excluded a small number of practices without a QOF score, as many of these practices are no longer active and we reasoned that any practice not participating in QOF would be less representative of a ‘typical’ GP practice. This may have excluded a small number of practices that opened since the 2015/16 QOF scores were calculated; however there are no grounds to believe that such practices would have been systematically different to the rest of our population with respect to low-value prescribing or factors associated with it. Due to a large sample size and large effect sizes, we obtained a high level of statistical significance in many of the associations we observed.

### Policy implications and interpretation

There are important implications to be considered in terms of making cost savings in prescribing. Co-proxamol, liothyronine and trimipramine illustrate a concerning phenomenon, where despite successful efforts to limit prescribing numbers, costs have risen sharply. For example, co-proxamol is expensive as it was removed from the drug tariff, meaning that any prescriptions for it have to be sourced as a ‘special’ order ^13^. There is limited regulation of the cost of such special orders, making real world cost savings on such drugs difficult until there are a very small number of total prescriptions.

The prescribing rate of liothyronine has been comparatively steady in recent years, though less than half practices prescribe it. However, the price has increased dramatically since the drug began being sold as a generic. It was noted in 2015 that the price had increased 40-fold to £152 for 28 tablets^14^; however our latest data show this to have further increased to £218 per 28 pills (or £412 per mean prescription). The reason for this is likely to be lack of sufficient competition in the market, with its only UK manufacturer having been subject to investigation by the UK competition commission^15^.

It is important to note that there is some variability in agreement on which medicines within this list should be considered low-value. For example, it is widely agreed that the evidence for the efficacy of glucosamine is limited^16^, and it is not recommended by NICE^2^. Liothyronine, conversely, has been restricted largely due to high cost rather than lack of efficacy. Similarly, gluten-free food prescribing is a controversial issue: guidelines and clinicians broadly agree that prescribing staple gluten-free foods such as bread and flour to patients with coeliac disease aids in adherence to a gluten-free diet; but prescribing has recently declined due to cost-saving measures initiated by NHS England and mediated by CCG policies.

Lastly, we are aware of no previous work showing that CCG policy has an impact on prescribing. Many maps in Appendix A show a large degree of variation according to CCG: while some variation may be due to demographic differences, it seems likely that much is due to variation in CCG policy. Substantiating this, CCG was significantly associated with prescribing behaviour in our mixed effects model. For medicines with limited efficacy, this merely represents CCGs varying in how quickly they can reduce costs. However, for efficacious treatments like gluten-free foods and liothyronine, such policy variation represents inequality of access to medication.

### Conclusions

The detailed analysis of trends and variation in low-value prescribing presented here should be used to inform future debate about the use of such medicines. A certain proportion of current variation in many of the observed medications is likely to be due to differences in policy between different CCGs and practices, rather than clinical need.

## Ethical approval

This study uses exclusively open, publicly available data, therefore no ethical approval was required.

## Guarantor

BG is guarantor.

## Contributorship

AW and BG conceived and designed the study. AW and HC collected and analysed the data with methodological and interpretation input from RC, SB and BG. AW drafted the manuscript. All authors contributed to and approved the final manuscript. SB was lead engineer on the associated website resource with input from RC, AW, BG and HC.

## Acknowledgements

Lead engineer on the original OpenPrescribing tool was Anna Powell-Smith.

